# The Extremely Brilliant Brain: An Isotropic Microscale Human Brain Dataset

**DOI:** 10.64898/2026.01.27.702076

**Authors:** Matthieu Chourrout, Andrew Keenlyside, Eric Wanjau, Yael Balbastre, Ekin Yagis, Joseph Brunet, David Stansby, Klaus Engel, Xiaoyun Gui, Julia Thönnißen, Timo Dickscheid, Laurent Lamalle, Alexandre Bellier, Umesh Vivekananda, Paul Tafforeau, Peter D. Lee, Claire L. Walsh

## Abstract

We present an isotropic 7.72 µm*/*voxel post-mortem human brain dataset acquired using Hierarchical Phase-Contrast Tomography (HiP-CT) at the ESRF Extremely Brilliant Source, beamline BM18. This fills a critical gap between whole-brain MRI at 100 µm resolution and serial-section histological reconstructions at 20 µm or finer. HiP-CT contrast, derived from X-ray phase shifts, enables rich 3D visualisation of complex neuroanatomy including white-matter bundles, microvasculature, and sub-nuclei. We provide open-source workflows for online data exploration, subvolume download, segmentation, and reintegration of analyses into the full dataset. We demonstrate the potential of this resource by tracing vasculature over long distances, segmenting nuclei, and extracting whitematter orientations with 3D structure-tensor analysis. High-resolution human brain datasets are transformative for quantitative neuroanatomy, circuit mapping, and validation of clinical imaging; this openly available resource is a critical step for global access to next-generation multiscale brain imaging.

---

High-quality and high-resolution reference datasets have enabled much of the research into multi-scale neuroanatomy, including cortical mapping, circuit tracing, and vascular mapping as well as providing validation for novel imaging methodologies. *In vivo* magnetic resonance imaging (MRI) reference datasets have reached an in-plane resolution of 0.19 mm × 0.19 mm [1], or an isotropic resolution of 0.25 mm [2], but are limited by motion-sensitivity and low signal-to-noise (SNR) regime. *Ex vivo* MRI retains this information-rich contrast while alleviating motion-related artefacts and increasing SNR *via* longer scan times, allowing an entire human brain to be reconstructed at 0.1 mm isotropic resolution after 100 hours of scanning [3].

Despite these advances, smaller structures in the brain still remain unresolvable with whole-brain MRI. For instance, individual cortical layers can have a thickness of 100–500 µm [4], while microvasculature diameters range from 15–240 µm (arterioles), 10–125 µm (venules) [5] down to 10 µm (capillaries) [6]. In the hippocampus, the granule cell layer of the dentate gyrus can have a thickness of 60 µm [7], all of which are well below the resolution of the most detailed MRI.

Microscopy sections provide additional information such as cell types, at the expense of three-dimensional (3D) consistency. When paired with serial sectioning and histological staining over an entire *ex vivo* brain, the data can be 3D reconstructed. The BigBrain dataset [8] was the first open-access histology-based 3D volume, with an isotropic resolution of 20 µm. This laborious acquisition process can span up to 1000 h [8], and requires extensive post-processing, barring its use even in small cohort studies.

Synchrotron X-ray phase-contrast tomography provides refractive index-based contrast for *ex vivo* soft tissues. Numerous studies [9–13] have used this technique to perform virtual histology in rodent brains and excised chunks of human brain parenchyma down to 2.54 µm*/*voxel in large slabs [14]. These methods present challenges when scaling to whole human brain due to sample preparation and data handling (an intact human brain at 2.54 µm*/*voxel would generate petabyte-scale datasets). In most cases, tissue preparation and scanning for synchrotron X-ray phasecontrast tomography involves fixation with formalin, dehydration through ethanol and immersion in ethanol during scanning [15] — this is sometimes followed by paraffin embedding to prevent motion of the sample during acquisition [16]. This type of sample preparation is much less aggressive than protocols used in light-sheet fluorescence microscopy, which ensures that the integrity of anatomical structures is preserved. The retrieved phase-contrast can be readily compared to either histology at high resolution or to structural MRI at lower resolutions.

Hierarchical phase-contrast tomography (HiP-CT) overcomes many of these limitations and enables scalable synchrotron X-ray tomography to whole human organs [17]. HiP-CT can be performed at the BM18 beamline of ESRF (The European Synchrotron, France) due to the fourth-generation upgrade of the X-ray source — the Extremely Brilliant Source (EBS) [18]. In the present work, scanning a complete adult human brain with HiP-CT required only 23 h for an isotropic voxel size of 7.72 µm. Motion and bubbling within the whole sample were prevented by mounting the sample in an agar/ethanol gel and by degassing [17].

*Ex vivo* imaging has inherent limitations. Tissue can degrade during the postmortem interval, and removal from the skull can damage delicate structures. For example, a recently discovered fourth meningeal layer was only visible using *in situ* imaging [19]. Fixation and dehydration also introduce changes: including shrinkage, and some distortion of soft tissue [20], especially in white matter. Despite these limitations, biobanks across the world have been collecting *postmortem* specimens for more than two decades, as fixed or frozen tissue of different sizes—from whole organs to tissue sections or biopsies.

Here we introduce the “Extremely Brilliant Brain” (EBB) dataset as a proof of concept for efficient microscale isotropic imaging of the intact human brain. Functioning as a digital biobank sourced from the Human Organ Atlas Hub, it enables long-term preservation and open access to microscale anatomical information that can be analyzed and integrated with complementary modalities. The release of this dataset, paired with *ex vivo* MRI and registered to the BigBrain space [8], together with examples of multiscale analyses, offers a new resource for the neuroscience community to perform virtual histology and to investigate microstructural features in minimally altered tissue.

## Results

The EBB dataset (Fig. 1) of a whole adult human brain acquired with HiP-CT contains 18 709 × 18 709 × 21 517 voxels with an isotropic voxel size of 7.72 µm, amounting to 13.7 TiB of data before masking and compression. The donor, pseudonymized as LADAF-2021-17, was a male who died of a pancreatic cancer at 63 years old. Based on the donor’s medical history and autopsy detecting no intracranial pathology or metastasis — apart from a small brain lesion (Fig. A1) —, this brain can be treated as a healthy specimen. After mounting using the previously described protocol [21], the brain measured approximately 15.5 cm along the anteroposterior axis, 12 cm along the mediolateral axis, and 11 cm along the dorsoventral axis. Prior to HiP-CT, it was imaged on a 3T MRI scanner at the IRMaGe MRI facility with a sagittal 3D TSE *T*_2_-weighted anatomical sequence to account for the ethanol-based mounting medium. The MRI sequence parameters are detailed in the Online Methods. The MRI scan and the HiP-CT dataset were rigidly registered. The HiP-CT setup and reconstruction pipeline are detailed in the Online Methods and in Fig. A2.

**Fig. 1.**
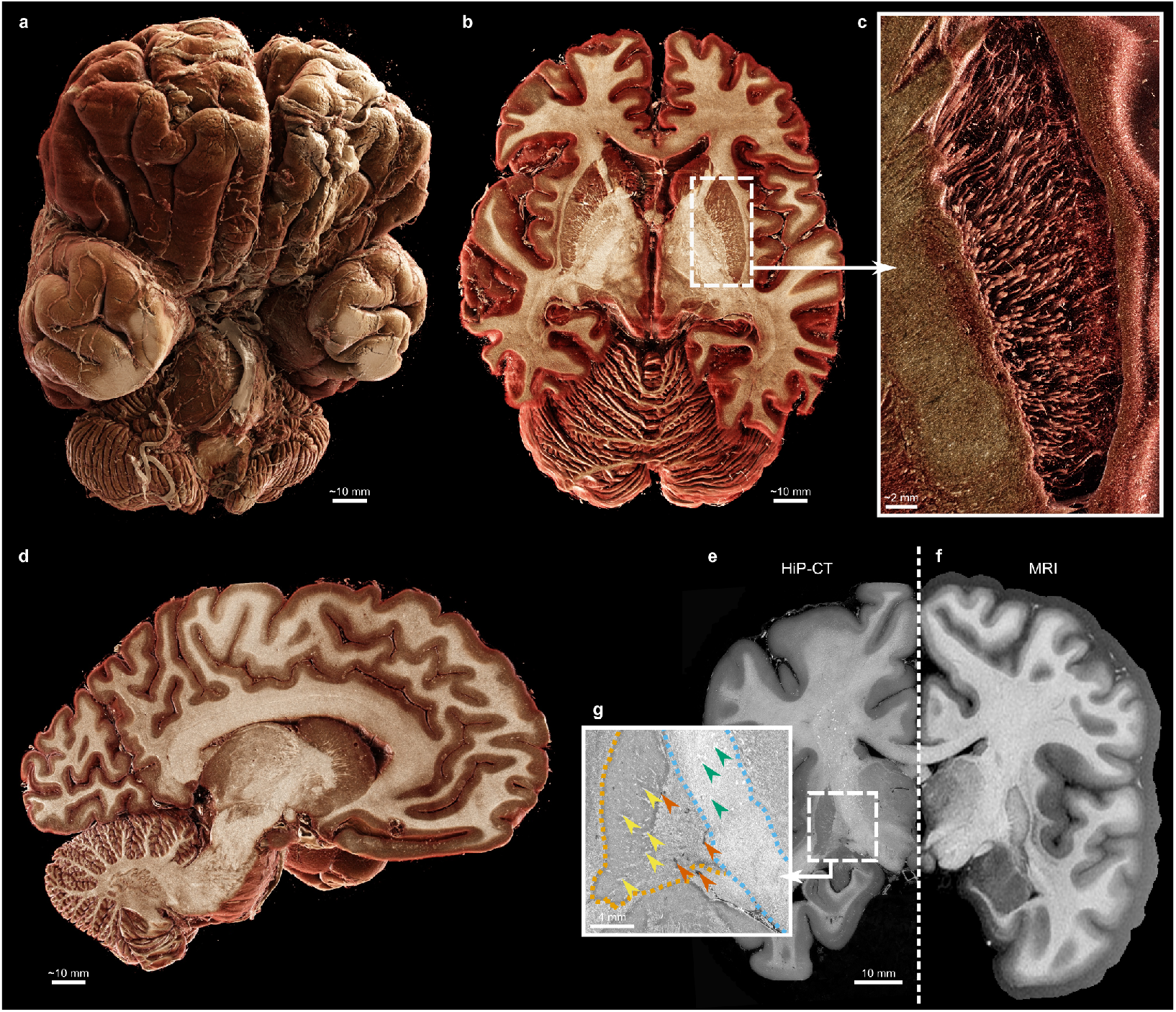
Overview of the HiP-CT “Extremely Brilliant Brain” (EBB) and of its paired MRI. **a-d**, 3D anatomical renderings of the brain, with an axial section (**b**), an inset (**c**) where white matter fibers of the left putamen are rendered across a few millimeters and a sagittal section (**d**). Renderings were generated using the Siemens Healthineers Cinematic Rendering from a binned-by-8 version (voxel size of 61.76 µm, *cf*. Table A2). **e**-**g**, Coronal slice of the “Extremely Brilliant Brain” dataset (**e**) and of its paired 3 T *T*_2_-weighted MRI (**f**) with an inset in the axial section (**g**) in the regions of the internal capsule (blue dashes) and lentiform nucleus (orange dashes) where the CLAHE filter emphasizes the region boundaries and the white-matter fibers (yellow arrowheads); blood vessels (orange arrowheads) and crystalline structures (green arrowheads) can also be spotted. The grayscale of the *T*_2_-weighted MRI was inverted to match HiP-CT contrast. A more detailed comparison between MRI and HiP-CT is provided in Fig. A4.

### Dataset accessibility

The complete dataset is available for download from the BioImage Archive (https://www.ebi.ac.uk/biostudies/bioimages/studies/S-BIAD1939) or from the data portal of the Human Organ Atlas (HOA), the wider HiP-CT organ imaging project (human-organ-atlas.ersf.eu). The latter also features other brains and other organs imaged using HiP-CT at a lower voxel size (around 20 µm per cubic voxel); volumes of interest within each of these organs at up to 1.3 µm*/*voxel are also available. All data can be browsed, downloaded and re-used under a Creative Commons Attribution 4.0 International license (CC BY 4.0).

The EBB dataset is provided in OME-Zarr [22] and JPEG2000 formats. The MRI dataset is provided as a compressed NIfTI. Metadata are provided with both the HiPCT and MRI data in a BIDS-compliant (https://bids.neuroimaging.io/) format—as TSV and JSON files. These standardized open formats are meant to ease the reuse of the data. Finally, the associated hoa-tools Python package (https://github.com/HumanOrganAtlas/hoa-tools) can interact with the EBB dataset to build pipelines in the Python programming language.

Due to the large size of the EBB dataset, few researchers will have the infrastructure or hardware to download and interact with the full-scale data, thus the data are made available for interactive exploration through a web-based viewer, Neuroglancer (Google, USA). To enhance reproducibility, usability and transparency, we provide interactive browser-based links for every figure in the manuscript, allowing the figure content to be examined dynamically within the viewer.

To align and facilitate comparison within the neuroscience community, the data is also released via EBRAINS (ebrains.eu), the European ecosystem for the brain research community, and is aligned in the same space as the BigBrain [8] (Fig. 2).

**Fig. 2.**
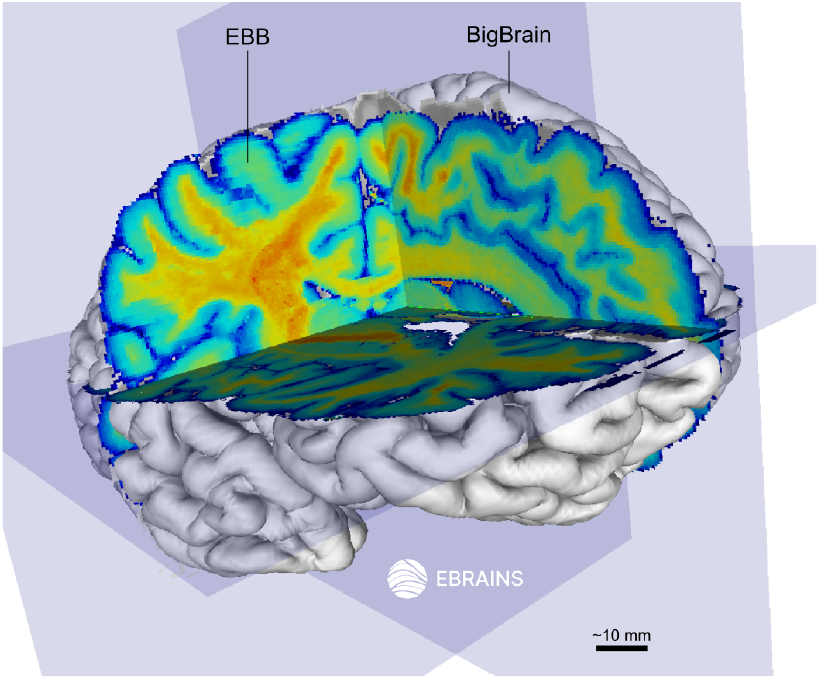
Integration of the “Extremely Brilliant Brain” (EBB) into the BigBrain space using the Siibra viewer (EBRAINS). The EBB cross-sections appear as yellow–red (white matter) and as green–blue (gray matter) in the BigBrain surface which is 3D-rendered in white.

We demonstrate how the release of the EBB dataset can foster neuroscience research with three examples of analysis. Our analyses target the vascular network, the hippocampus and the white matter tracts, which are involved in neuropathologies such as stroke, dementia and epilepsy. We used WEBKNOSSOS (Scalable Minds, Germany) for web-based annotation and collaborative multi-scale segmentation of the blood vessels; FastSurfer [23] for whole-brain segmentation of the cortical structures and nuclei; HippUnfold [24] for parcellation of the hippocampus in the paired MRI.

### Cloud-based blood vessel segmentation

To illustrate the ability to collaboratively interact with and manually segment or quality control automatically segmented features in the data we segment a meningeal vessel, using large-scale data browsers and optimised cloud storage data formats. A large meningeal vessel was labelled through the brain, along with its major branches over 8 cm. The lumen of the vessel was segmented in 3D from the inferior to the superior brain surface as it travelled in the longitudinal fissure over a distance of 8 cm. The vessel network begins at the left anterior cerebral artery (Fig. 3, black arrow) along with its A1 (Fig. 3.1) and A2 segments (Fig. 3.2), which later becomes the callosomarginal artery (Fig. 3.3) then the paracentral artery (Fig. 3.4) as it reaches the left paracentral cortex up to the central sulcus [25]. Although there are a few discontinuities in the segmentation due to the collapse of the vessel wall, this long-range tracking highlights feasibility for vascular mapping across multiple length scales.

**Fig. 3.**
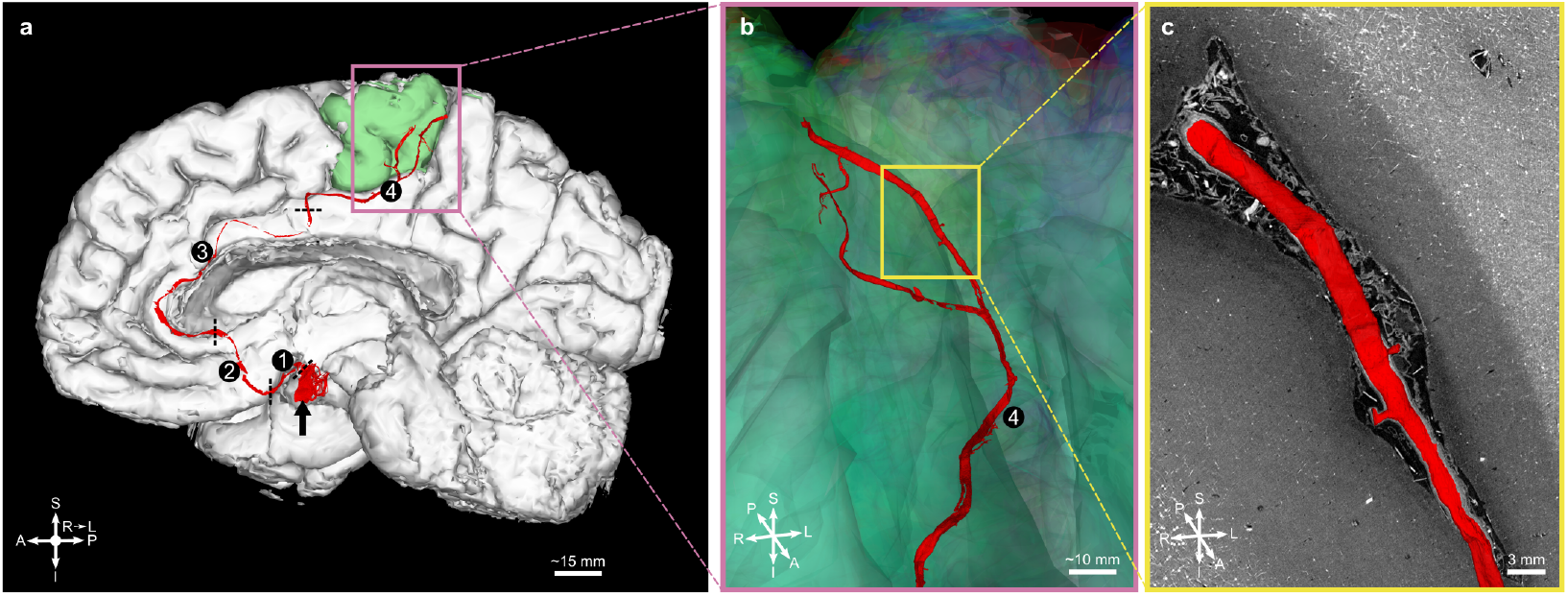
Segmentation of the anterior cerebral artery and of its branches. **a**, Anterior cerebral artery (black arrow) along its segments A1 ➀ and A2 ➁ and its distal branches — the callosomarginal artery ➂ and the paracentral artery ➃ — up to the paracentral cortex (green) from the FastSurfer output. **b**, Close-up view in a different perspective. **c**, 2D cross-section with the 3D rendering. S indicates superior; I, inferior; A, anterior; P, posterior; R, right; L, left.

**Fig. 4.**
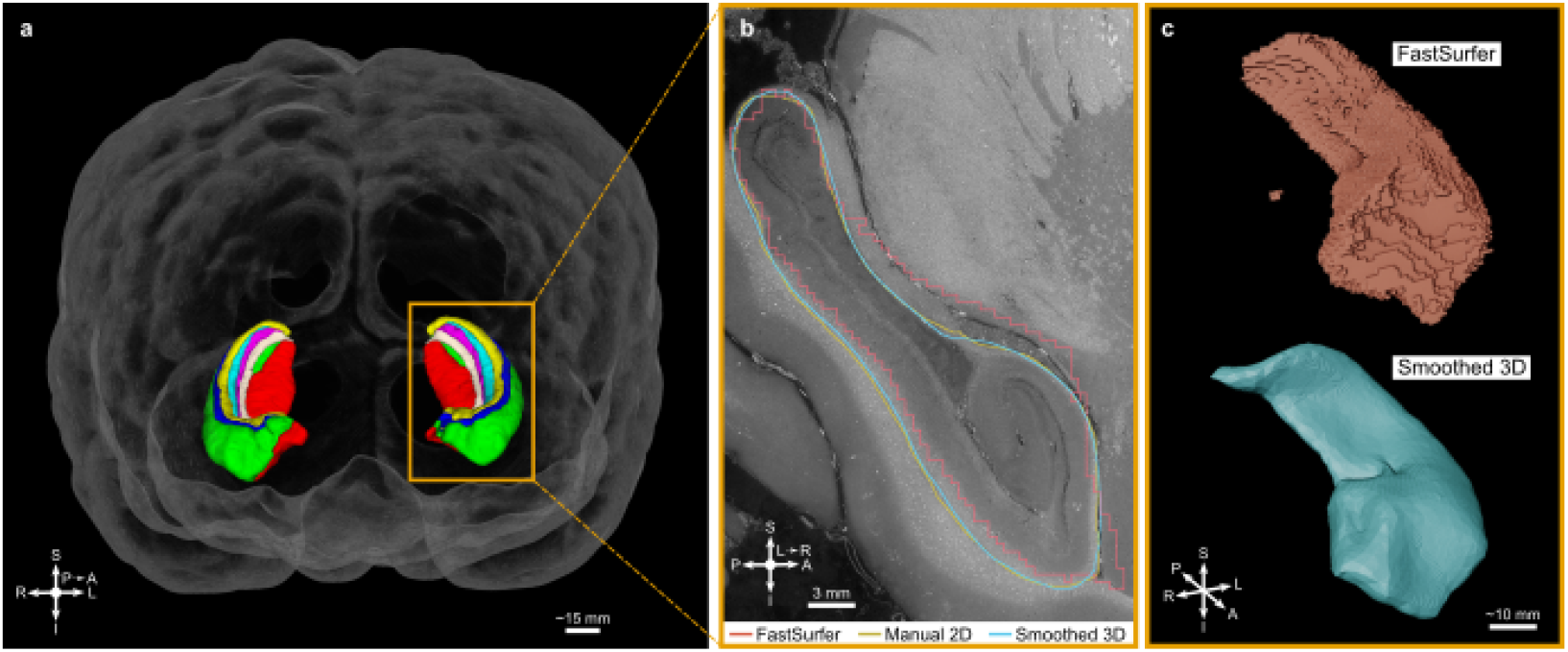
Segmentation of the hippocampi. **a**, Hippocampal subfields determined from the paired MRI using HippUnfold (Fig. A5). **b**,**c**, Comparison between the initial label of the left hippocampus from FastSurfer (Fig. A6), the refined manual segmentation and the final smoothed segmentation along a sagittal slice (**b**) and as 3D renderings (**c**). S indicates superior; I, inferior; A, anterior; P, posterior; R, right; L, left.

### Parcellation of hippocampi

As HiP-CT contrast appears similar to a *T*_1_-weighted contrast in MRI, many tools developed by the MRI community can be readily applied to downsampled HiP-CT data to provide fast initial segmentations and parcellations. These can then be refined using the far higher resolution of the HiP-CT, which thus greatly speeds up analyses. The hippocampus was initially segmented using currently available MRI tools: HippUnfold [24] on the MRI dataset and FastSurfer [23] on the EBB dataset; for the latter, the EBB dataset was binned to 494.08 µm. Using our interactive online segmentation tools, the surface of the hippocampus (Fig. 4) was re-contoured with neuroanatomy expertise on the full 7.72 µm data. The Dice similarity coefficient between the FastSurfer surface and the new surface was computed for each pair of manually annotated slices (mean of 0.595) and on the final volume after smoothing (mean of 0.627). HiP-CT enables contrast and resolution of hippocampal layers in 3D (Fig. 4b) that cannot be seen in MRI and would allow for a more precise delineation of the layers and structures.

### Predominant orientations of white matter fibers

White matter in the brain exhibits a multi-level hierarchical organization where individual axons bundle together to form larger fiber tracts (Fig. 1d). White matter fibers are visible in the EBB as the surrounding myelin provides a sufficient phase shift to be detected by synchrotron phase-contrast [15].

Gradient-based structure tensor analysis (eigen decomposition of image intensity gradients) [26] can retrieve the predominant orientation of these white matter fibers, as shown in Fig. 5. Computation was performed on a 0.5-mm-thick coronal slab, which was efficiently retrieved from the whole dataset thanks to its chunk-based storage layout. Parameters (*cf*. Online Methods) were empirically chosen after testing multiple configurations to balance sensitivity and specificity of fiber orientation detection.

**Fig. 5.**
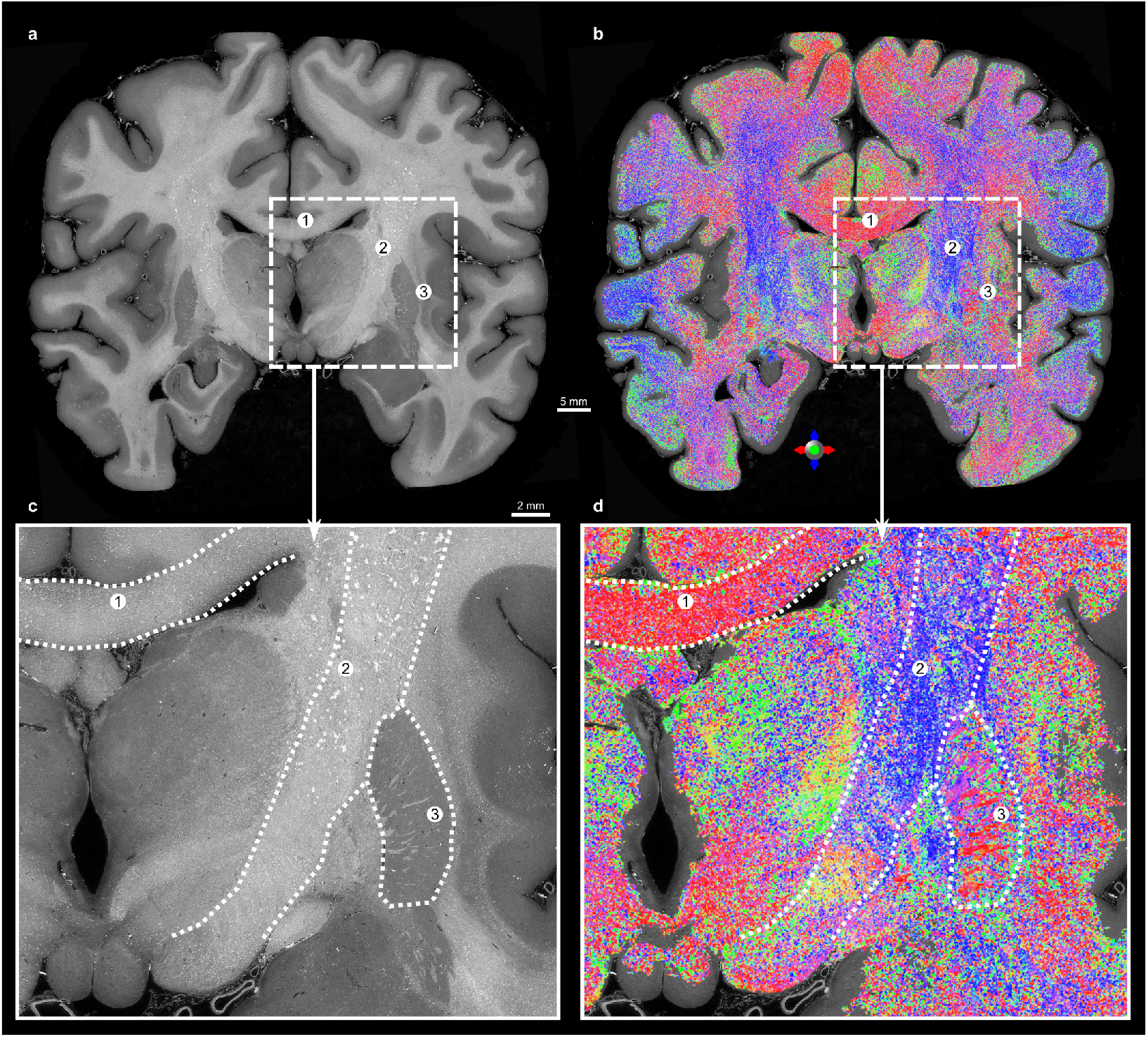
Orientation of the white matter fibers can be retrieved with structure tensor analysis. **a**, Coronal section of the “Extremely Brilliant Brain”. **b**, Direction-encoded colors (DEC) derived from the structure tensor analysis over a 64-voxel-thick slab within a mask of white matter overlaid on the same section. **c**,**d**, Insets with views in the HiP-CT coronal section (**c**) and in the DEC map (**d**). Regions described in the main text: ➀ lentiform nucleus. the corpus callosum, ➁ the corticospinal tract and ➂ the lentiform nucleus.

In the corpus callosum (Fig. 5.1) and in the corticospinal tract (Fig. 5.2), the retrieved orientations are respectively along the left–right and superior–inferior axes, as expected [27]. Fibers in the lentiform nucleus (Fig. 5.3) can be seen projecting laterally across the putamen, against the gray matter. The fibers in these regions cannot be distinguished in the ∼255 µm paired MRI, but they have already been visualised in a 100 µm *ex vivo* MRI dataset [3] (Fig. A4).

The ability to compute the structure tensor over a whole human brain coronal section at the native voxel size (7.72 µm) is a first step towards high-resolution white matter characterisation.

## Discussion

The EBB dataset represents an important addition to the whole human brain atlas community. Spanning the entire brain with an isotropic voxel size of 7.72 µm and X-ray phase contrast provides many new opportunities to study multi-scale brain connectivity and cytoarchitecture, and to link to existing brain atlases from complementary imaging sources such as the BigBrain [8]. HiP-CT contrast derives from the phase shift of X rays travelling through the tissue, providing contrast for a large range of structures in a label-free manner. Given the size of the EBB dataset, many features and structures have not been analysed, and full assessment of the data is beyond the capacity of any single research group. Thus the key challenge to solve was to design end-to-end pipelines that enable efficient interaction with the global community and test these pipelines through example analyses with collaborators, in neuro-anatomy and clinical neuroscientists as well as image analysts. This required careful selection of data formats and processing tools. We generated multiple binned versions of the datasets, enabling tailored use (Table A2). This strategy allowed the dataset to be efficiently integrated with common neuro-imaging pipelines.

We first showcased the segmentation of a blood vessel over 8 cm (Fig. 3), which was created by seeding and growing regions in the lumen on the 7.72 µm data. In *ex vivo* imaging, like HiP-CT, vessels have a tendency to collapse during sample preparation [28]. As the lumen can completely disappear when the vessels fully collapse, focusing on the walls of the arteries, which are hyperintense in HiP-CT, is an alternative approach to segment parts of the vascular network.

The hippocampal layers are well delineated in HiP-CT, which also benefits from a higher resolution than MRI. Thus, the labels of the outer surface of the left hippocampus were refined under the supervision of neuro-anatomy experts using the cloud-based WEBKNOSSOS tool as a collaborative platform. Starting from the parcellation generated by an automated tool (FastSurfer, in this case) substantially reduced manual effort and enabled comparison with MRI-like resolution. This dataset has enabled the precise segmentation of the left hippocampus in a 3 cm × 3 cm × 4 cm region of interest. Extracting white matter fiber orientations from the image gradient has been applied to multiple microscopy modalities. With histology, it can be applied to the 2D images from the tissue sections to retrieve the 2D orientation [29]. Histology-derived orientation has also been successfully correlated to the paired diffusion MRI [30]. While a range of softwares have been developed to compute the structure tensor, datasets of the EBB’s size require bespoke implementations that are developed with scalability and compatible with chunk-based image data formats [31]. A few pioneer studies have been able to retrieve tractography from synchrotron tomographic data [32, 33]. Because of the higher voxel size, such tractography could resolve ambiguities in diffusion MRI, including fiber crossings and branchings [34]. This advance would refine the mapping of tracts within nuclei such as the zona incerta and subthalamic nucleus, which are key targets for deep brain stimulation in epilepsy and Parkinson’s disease [35], thereby improving surgical precision and patient outcomes. However, as shown in Fig. 5, such approaches requires choosing parameters that favour a particular object size. Besides, confounding elements might alter the structure tensor; these can be blood vessels (similar rod-like shape as fibers), noise (due to either acquisition or unresolved details), or hyperdensities (either pathological [36] or exogenous).

By releasing the EBB and its paired MRI, we are enabling a wide range of researchers to access and work with this data. We have presented some example analysis for vasculature, hippocampal parcellation and white matter bundles, but there are many others that could be explored by the global community.

Several studies have mapped smaller brain regions using high-resolution techniques or devices optimized for limited samples sizes [37, 38]. HiP-CT serves as an effective intermediate imaging modality, bridging the resolution gap between macroscopic imaging techniques (*>* 0.5 mm, e.g. MRI, CT) and microscopic techniques (*<* 1 µm, e.g. optical microscopy, EM [39]). As HiP-CT does not damage the tissue, applying HiP-CT to biobanks could provide a wealth of high-resolution images to supplement the targeting of further destructive testing or detection of cohorts of pathology not visible on conventional imaging, without limiting the choice for future studies, as the tissue can be returned to the biobank after HiP-CT.

Although the EBB dataset does not resolve cellular structures, it could provide a more detailed insight into white matter organization, in a similar fashion to NODDI or diffusion MRI metrics, which are of clinical relevance for a wide range of neuropathologies, including Alzheimer’s disease, Parkinson’s disease, and multiple sclerosis [40].

Atlases of the pial arteries are usually created with time-of-flight MRI scans at the resolution of the parcellation atlases (in-plane resolution: 0.5 mm; slice thickness: 0.8 mm) [41]. Using the EBB to jointly develop a high-resolution 3D vasculature segmentation and a high-resolution white matter mapping would help to understand the interaction between the axonal and the vascular networks. This could improve stroke management by enabling the recognition of spatial patterns associated with clots or aneurysms and by helping to identify collateral blood supply relevant to recovery.

Although the EBB provides a highly singular resource, there are known imperfections with the dataset stemming from sample prep and scanning which can and have been improved upon since the collection of the EBB dataset [42]. The EBB exhibits some shrinkage compared to an *in vivo* brain due to the formalin fixation and dehydration; also a number of bubbles formed during previous scans of the sample (which are also available on the HOA data portal) due to the radiation [43]; these bubbles are mainly located in the right-hemisphere striatum and in the ventricles, and they create streak artifacts during reconstruction. Despite these imperfections the EBB still provides a high-resolution 3D dataset without mechanical sectioning, preserving structural integrity and without rips or tears which usually appear with serial sectioning. In addition, as HiP-CT is non-destructive, the brain is still available and allows for further scans of the same sample in the future as the technology develops and matures [42].

The EBB was collected over 23hrs, a relatively fast data acquisition, which shows clear promise for larger cohort studies. The analyses demonstrated here could potentially be performed and compared across multiple individuals and would provide insight into structural variability in healthy and path-alogical context.

Vessel segmentation could be applied as method for understanding variation the aetiology of vascular disorders in the brain. The enlarged perivascular spaces [44] are a shared biomarker among cerebrovascular diseases that could be studied with synchrotron imaging. More specific biomarkers could be screened in the blood vessel wall, although it is still unclear whether they would be revealed by synchrotron imaging; for instance, cerebral amyloid angiopathy is characterized by deposits of amyloid protein [45], whereas atherosclerosis involves the accumulation of lipid-rich plaques within the arterial intima [46]. A Kaggle competition has previously been held for AI-based vasculature segmentation in HiP-CT kidney data [47], and there is ongoing work on fine-tuning the highest-ranked models towards HiP-CT brain data.

Clinically, subfield-by-subfield quantification improves early detection and stratification of cognitive decline thereby supporting individualized therapeutic decision-making [48]. In medial temporal lobe epilepsy, precise delineation of hippocampal arterial complexes and their relationships to the anterior choroidal artery and the posterior cerebral artery helps plan safer resections (with preservation of high-risk perforators) while maximizing sparing of memory-related circuits [49]. Finally, vasculo-anatomical segmentation enables estimation of hippocampal vascular reserve indices, which are associated with cognitive performance and hippocampal volume, opening the way to precision medicine that integrates structure, vascularization, and cognition [50]. Additionally, HiP-CT vascular geometries offer high-fidelity templates to initialize and validate donor-specific hemodynamic simulations, thereby supporting risk stratification and surgical planning across cerebral territories, including the AComA/ACA complex [51]. Finally, by mapping perforators and collateral pathways at branch-level resolution, HiP-CT can inform interpretation of stroke phenotypes and the prognostic role of leptomeningeal collaterals measured *in vivo*, linking *ex vivo* anatomy to clinically actionable endpoints [52].

The release of the EBB together with the *ex vivo* MRI and curated segmentations provides a substantial and accessible resource for the neuroimaging community. In addition to offering new, openly available materials, it includes concrete examples and guidance for data-handling tools capable of operating at terabyte scale. This enables researchers to design and deploy pipelines that manage high-resolution volumes efficiently, lowering technical barriers and broadening community access. This work supports a deeper integration of synchrotron-based imaging into multimodal neuroanatomical studies. HiP-CT can now enrich the knowledge base at this micrometric scale, making previously inaccessible levels of structural detail available for systematic investigation. The potential impact spans both basic and clinical neuroscience: from refining disease models and translational frameworks to improving the resolution and composition of research cohorts.

## Supporting information

Supplemental Media 1

Supplemental Media 2

## Supplementary information

The following supplementary material is included:

- Cinematic Anatomy animations

## Acknowledgements

We would like to thank the donor and their family. We acknowledge the European Synchrotron Radiation Facility (ESRF) for provision of synchrotron radiation facilities, notably its beamline BM18 (proposals MD-1290, MD-1389). The IRMaGe MRI facility is partly funded by the French program “Investissement d’Avenir” run by the French National Research Agency, grant “Infrastructure d’avenir en Biologie Santé” (ANR-11-INBS-0006). This work was made possible by the facilities and support provided by the Research Complex at Harwell and Royal Academy of Engineering (CiET1819/10). This project has been made possible in part by grant number 2020-225394 from the Chan Zuckerberg Initiative DAF, an advised fund of Silicon Valley Community Foundation, and by grant number CZIF2021-006424 from the Chan Zuckerberg Initiative Foundation. Research reported in this publication was supported by the National Institute of Neurological Disorders and Stroke of the National Institutes of Health under award number UM1NS132358. The content is solely the responsibility of the authors and does not necessarily represent the official views of the National Institutes of Health. P.D.L. is a CIFAR Fellow in the CIFAR MacMillan Multiscale Human program.

## Declarations

### Ethics approval and consent to participate

A complete brain was acquired from Laboratoire d’Anatomie des Alpes Françaises (LADAF) in accordance with the French legislation on body donation. Written informed consent was obtained from the donor prior to death. All dissections were conducted with respect for the deceased’s memory.

### Consent for publication

Not applicable.

### Data availability

The “Extremely Brilliant Brain” dataset can be downloaded or browsed on the Human Organ Atlas data portal (https://human-organ-atlas.esrf.fr/), in the BioImage Archive (https://www.ebi.ac.uk/biostudies/bioimages/studies/S-BIAD1939) or on the EBRAINS page (https://search.kg.ebrains.eu/instances/fe09ed63-983f-4832-b7e3-e1131a95b870); its DOI is 10.15151/ESRF-DC-2268032801. Besides, the raw data is available under a Creative Commons Attribution 4.0 license at https://doi.org/10.15151/ESRF-ES-873257979.

### Materials availability

The interactive figures are provided in the GitHub repository (https://github.com/UCL-MXI-Bio/2025-chourrout-extremely-brilliant-brain), 12 the associated data is hosted in the BioImage Archive (https://www.ebi.ac.uk/biostudies/bioimages/studies/S-BIAD1939).

### Code availability

The code to convert data formats and interact with the data is provided in the GitHub repository (https://github.com/UCL-MXI-Bio/2025-chourrout-extremely-brilliant-brain).

### Author contribution (according to the CRediT taxonomy)

*Conceptualization* A.B., P.T., P.D.L., C.L.W.

*Data curation* M.C., A.K., J.B., D.S., L.M., P.T., C.L.W.

*Formal analysis* M.C., P.T., C.L.W.

*Funding acquisition* P.T., P.D.L., C.L.W.

*Investigation* M.C., A.K., E.W., E.Y., P.T., P.D.L., C.L.W.

*Methodology* J.B., T.D., L.M., P.T., P.D.L., C.L.W.

*Project administration* P.D.L., C.L.W.

*Resources* J.B., A.B.

*Software* M.C., J.B., D.S., X.G., J.T., P.T.

*Supervision* T.D., P.D.L., C.L.W.

*Validation* T.D., A.B., P.T., P.D.L., C.L.W.

*Visualization* M.C., A.K., J.B., K.E., A.B., P.T.

*Writing – original draft* M.C., Y.B., E.Y., J.B., U.V., A.B., C.L.W.

## Online Methods

### Sample preparation

A complete brain was acquired from Laboratoire d’Anatomie des Alpes Françaises (LADAF) in accordance with the French legislation on body donation; the donor (pseudonymised as LADAF-2021-17) was a 63-year-old male who died of pancreatic cancer. Written informed consent was obtained prior to death. All dissections were conducted with respect for the deceased’s memory.

The whole body was perfused with formalin via the right carotid artery at the time of death, followed by storage at a temperature of 3.6 ^°^C. The brain was post-fixed in 4 % neutral buffered formalin for a duration of 4 days at 3.6 ^°^C. Following fixation, the brain underwent stepwise dehydration through graded ethanol solutions up to 70 % concentration to minimize tissue distortion. The brain was then mounted in a container and embedded in an agar-ethanol gel for stabilization. To prevent the bubble formation during imaging, multiple thermal degassing cycles were applied at each stage of the dehydration and embedding processes. For more information about the preparation protocol, refer to [21].

### Data acquisition and reconstruction

The brain was first scanned with a 3 T MRI device (Philips Achieva 3.0T dStream, Philips Healthcare, Best, The Netherlands) at the IRMaGe MRI facility. The MRI sequence was a sagittal 3D TSE *T*_2_-weighted anatomical sequence (3D View Brain *T*_2_, 40^°^ constant refocusing control, TSE factor of 71, TR of 3000, TE of 280, 90^°^ flip angle, CS factor of 4.5, 200 mm × 200 mm × 150 mm (AP by HF by LR) field of view, 0.50 mm × 0.50 mm × 0.39 mm acquisition resolution, interpolated at reconstruction to 0.255 mm × 0.255 mm × 0.257 mm, 8 averages, 160 min acquisition duration).

The brain was then scanned using Hierarchical Phase-Contrast Tomography (HiP-CT) [17] at the BM18 beamline of the European Synchrotron Radiation Facility (ESRF), under proposal MD-1290. The distance between the X-ray source and the sample was 177 m, and the propagation distance was 10 m. To image the whole organ, we used a parallel polychromatic X-ray beam filtered through attenuators; the resulting average X-ray energy was 111 keV. The X-ray detector consisted of a LuAG:Ce 2000 µm reflective scintillator (custom-made by Crytur, Czechia; https://www.crytur.com/) combined with DZoom optics (ESRF, France) and an Iris 15 (Teledyne Photometrics, USA; https://www.photometrics.com/) camera (5056 px × 2960 px). To enhance the lateral field of view to ∼144 mm, a quarter-acquisition [43] was used — a first, central scan and a second, annular scan at the same vertical position are stitched together (prior to reconstruction, i.e. as projections) into one “stage”. The vertical field of view of each stage was 6 mm. The number of projections per stage was 30 000. An automated series was used with a vertical step of 5 mm between stages, corresponding to 33 stages with a 16.7 % overlap. The whole brain was imaged with a 3D isotropic voxel size of (7.72 µm)^3^ in 23 hours. A second sealed container, filled with ethanol 70 %, was scanned as reference for flat-field correction. All the stages for each angular position were stitched together in order to generate radiographic frames of the complete vertical size of the brain. An average projection of all these frames was performed and then filtered with an in-house processing tool in order to extract a low-frequency correction map that was then subtracted from all the complete frames in order to correct for the vertical inhomogeneities of the beam vertical profile, and to correct the largest ring artefacts. The complete radiographic frames were reconstructed into a volume of 13.70 TiB (or 15 TB) using a filtered back-projection algorithm coupled with a 2D unsharp mask filter (*c*_*unsharp*_ = 1.2, *σ*_*unsharp*_ = 4 px) applied on the complete frames and a single-distance phase retrieval 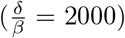, as implemented in the software package PyHST2 [53]. A synthesis of the scanning and reconstruction parameters can be found in Table A1.

### Registration into the BigBrain space

The Extremely Brilliant Brain was registered into the BigBrain space using voluba [54]. The transformation is affine and can be described as the following homogeneous matrix (lengths are in mm):

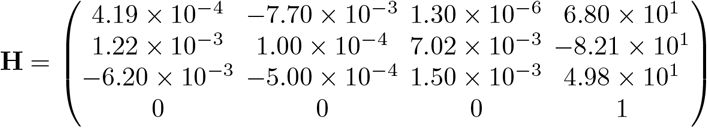

### Vessel study

The whole volume was loaded from the cloud storage into the official Webknossos instance (Scalable Minds). A large vessel was segmented using the online tools, especially the “Segment Anything 2”-powered Quick Select tool on the lumen of the blood vessel and the Interpolation tool. The segmented volume was downloaded from the official Webknossos instance then converted to a label (as a Neuroglancer precomputed volume).

### Parcellation of hippocampi

First, the background in the *T*_2_ MRI data was masked out manually using 3D Slicer (Brigham and Women’s Hospital, Boston, MA, USA & 3D Slicer contributors). Second, after conversion to NIfTI, the masked MRI data was processed using HippUnfold [24]. The generated parcellation was converted to the Neuroglancer precomputed format for rendering.

The EBB dataset was first downsampled to 494.08 µm*/*voxel and the background was masked out manually using 3D Slicer (Brigham and Women’s Hospital, Boston, MA, USA & 3D Slicer contributors). Second, after conversion to NIfTI, the brain was reoriented to match the standard axes using FreeSurfer (version 7.4.1; Massachusetts General Hospital, Boston, MA, USA). The whole brain was then segmented using FastSurfer [23] and the hippocampi were extracted as labels. These labels were loaded into the official instance of Webknossos (Scalable Minds, Germany) and the outer surface of the left hippocampus was manually redrawn at 7.72 µm*/*voxel using the FastSurfer output as a prior, driven by the HiP-CT image intensities and under the supervision of expert neuroanatomists; 200 slices were labelled over 4000 slices (one slice every 20 slices) then interpolated in 3D and smoothed using Amira3D (version 2022.2; Thermo Scientific, USA). The Dice similarity coefficient between the FastSurfer surface and the new surface was computed for each pair of manually annotated slices and on the final volume after smoothing. The generated segmentation was converted to the Neuroglancer precomputed format for rendering.

### Structure tensor analysis

The structure tensor was computed in a region of interest of the reconstructed data to retrieve the local orientation of the white-matter fibers using the cardiotensor Python package [31]. The computation relies on two convolutions, first by a Gaussian derivative kernel ∇_*σ*_ then by a Gaussian kernel *K*_*ρ*_ for integration [26]: **S** = *K*_*ρ*_ * (∇_*σ*_*V* (∇_*σ*_*V*)^*T*^); *σ* and *ρ* are the kernel sizes and they are adjusted depending on the size of the object of interest. The configuration file for cardiotensor is available to reproduce the vector field. The vector field was converted to an RGBA map (as a Neuroglancer precomputed volume) for rendering.

## Appendix A Extended Data

**Fig. A1.**
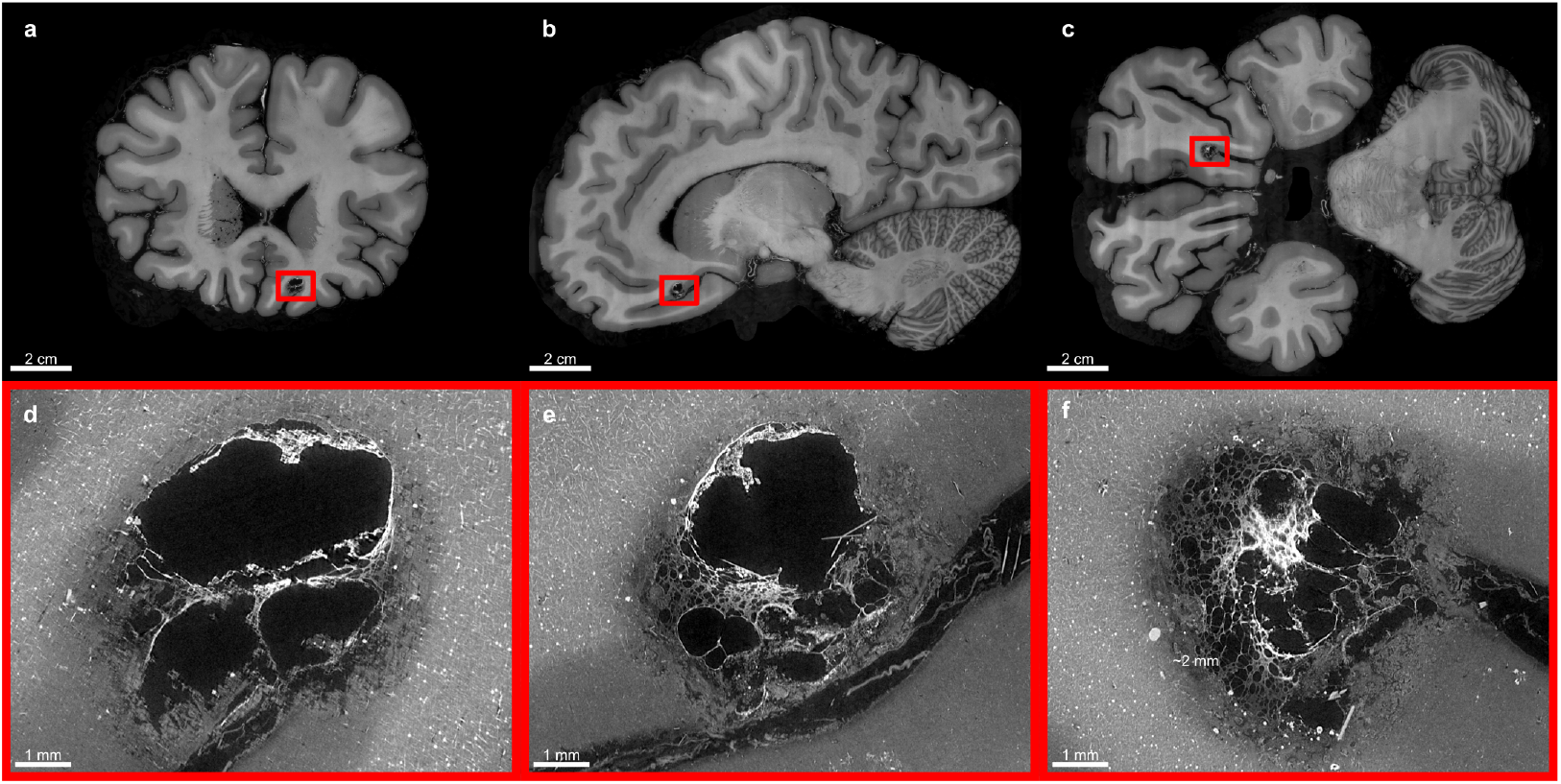
Brain lesion discovered with HiP-CT in the “Extremely Brilliant Brain” (EBB) dataset. **a, b, c**, Whole-brain views in the coronal plane (**a**), sagittal plane (**b**) and axial plane (**c**) to locate the lesion. **d, e, f**, Focus on the lesion in a small field of view in the coronal plane (**d**), sagittal plane (**e**) and axial plane (**f**). This could be either a cavernoma or a multiloculated pyogenic cortical brain abscess.

**Fig. A2.**
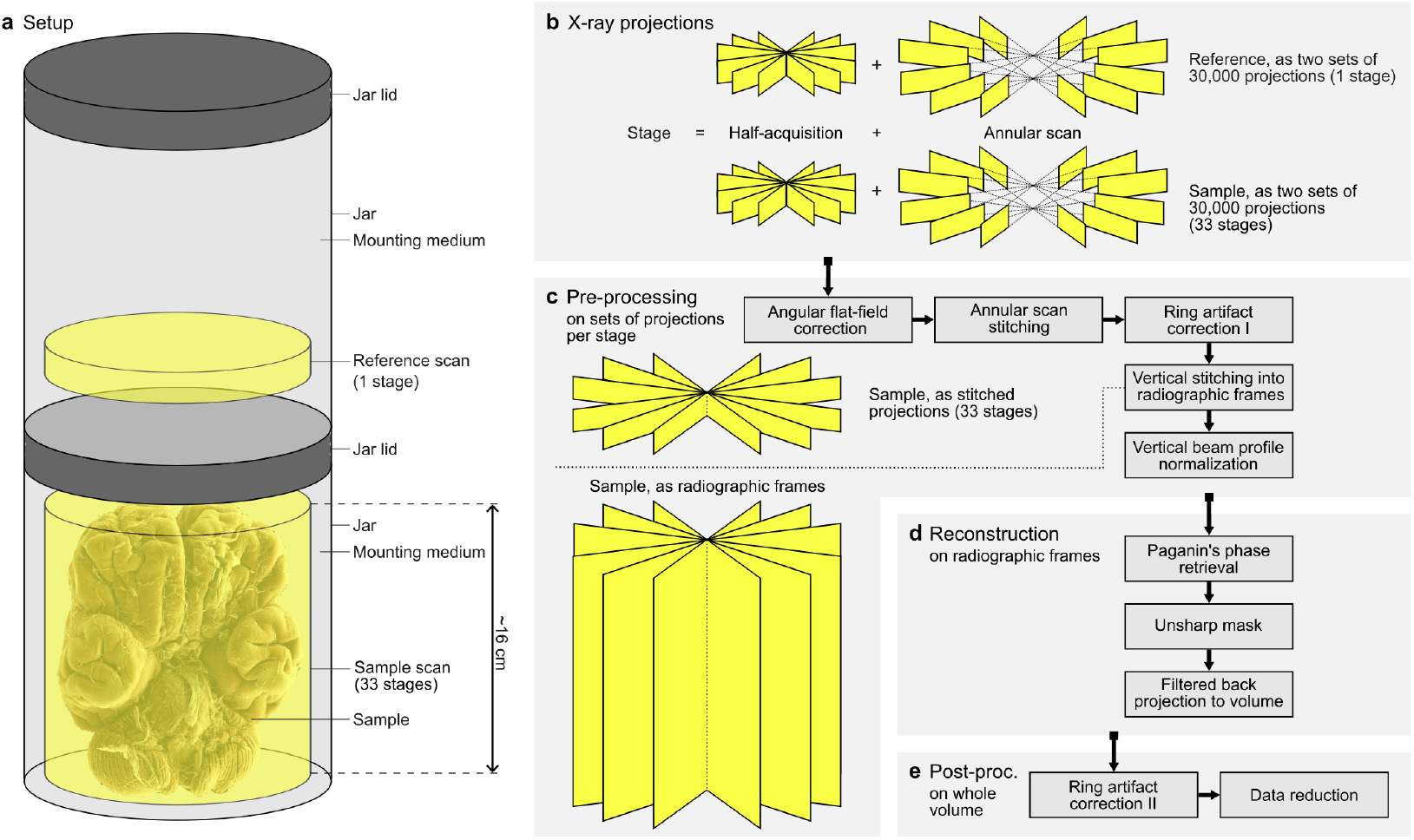
**a,b**, Scanning setup for HiP-CT. The HiP-CT setup (**a**) uses two identical containers: the sample is immersed in the mounting medium (agar/ethanol mix) [21] in the bottom jar while the top jar is filled with the mounting medium only to be used as a reference for the flat-field correction. In common synchrotron tomography setups, the sample is either immersed in phosphate-buffered saline or ethanol (or eventually embedded in paraffin) and the reference for the flat-field correction is acquired in the air. To extend the vertical field of view, multiple stages are acquired with an overlap. To extend the lateral field of view, a half-acquisition is combined with an annular scan (**b**); this is referred to as quarter-acquisition [43]. One stage is acquired in the reference, which is later used for the flat-field correction of all the stages which are acquired in the sample. **c**,**d**,**e**, Reconstruction pipeline for HiP-CT. As part of the pre-processing (**c**), the projections from the half-acquisition and from the annular scan are stitched together horizontally (per stage) then vertically (across stages). The reconstruction (**d**) extracts the phase shift in the stitched projections and creates a volume by filtered back projection. The post-processing (**e**) converts the numeric values to a usable range in the 16-bit unsigned integer format. Modified from [55].

**Fig. A3.**
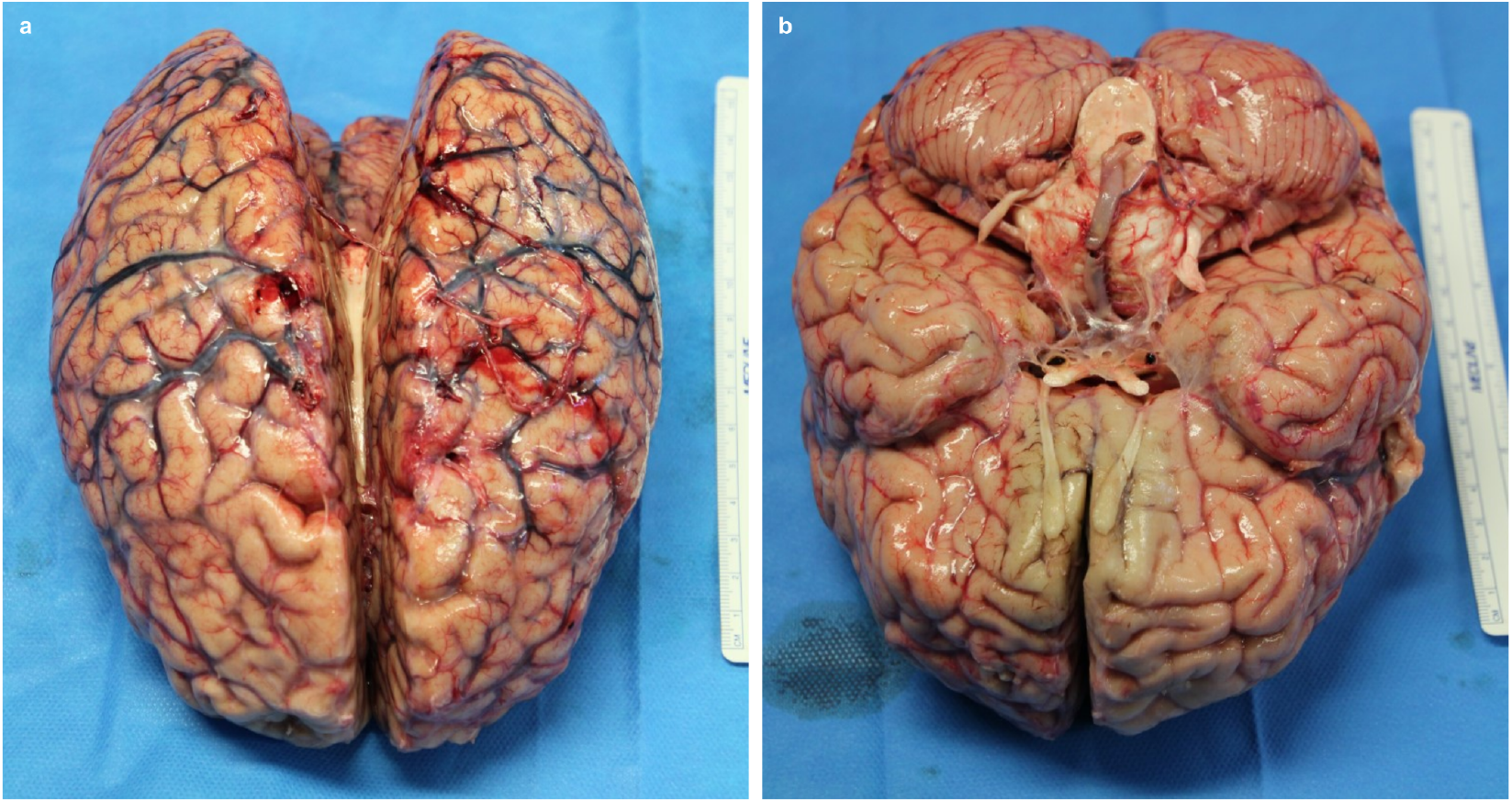
Dorsal (**a**) and ventral (**b**) photos of the whole brain post-autopsy. Although the body was perfused with formalin, blood was still present in the vascular network. The ruler on the side is 15 cm long.

**Fig. A4.**
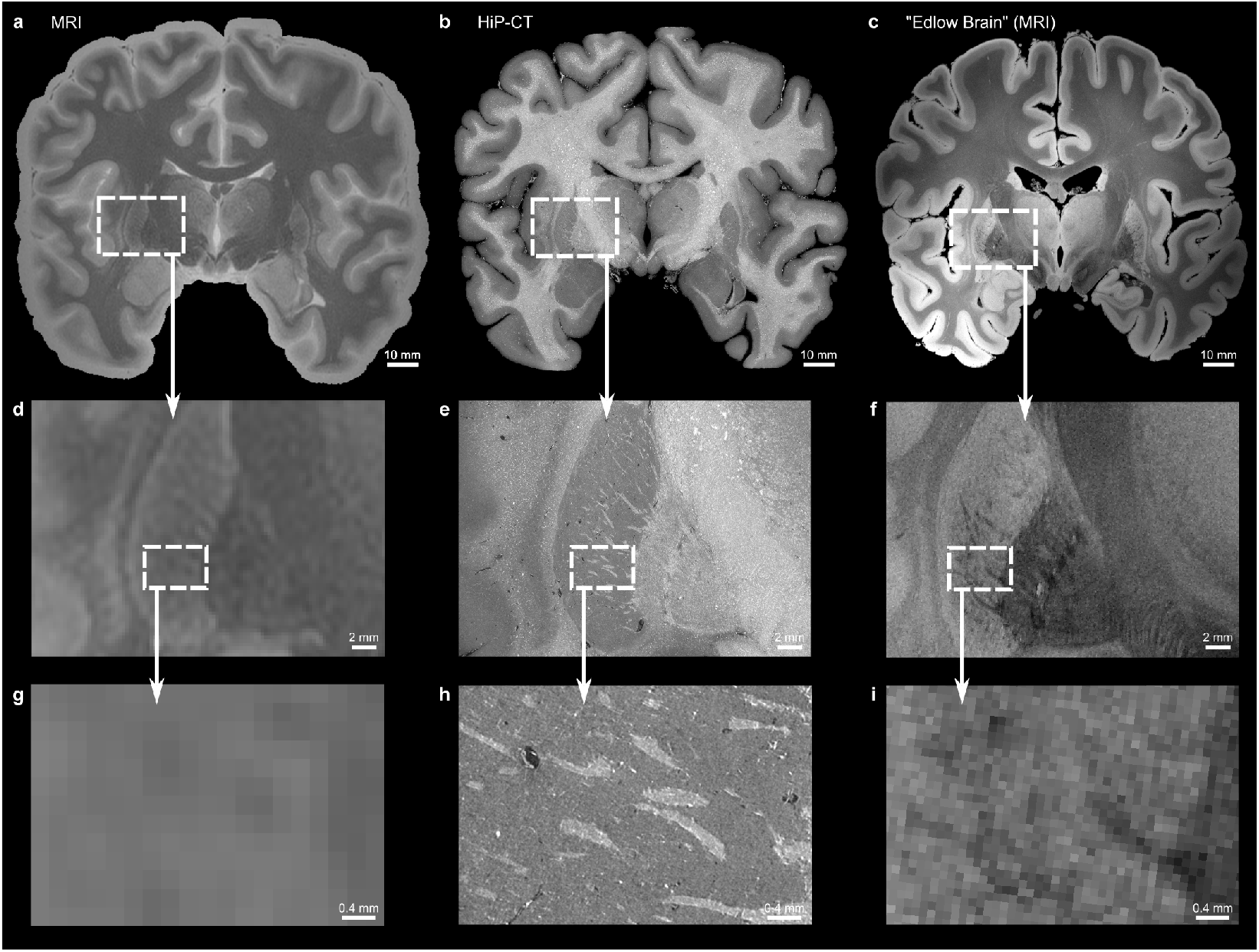
Comparison of the white matter fibers in the lentiform nucleus between the 3 T *T*_2_-weighted MRI (in-plane resolution: 0.255 mm; slice thickness: 0.257 mm) (**a**,**d**,**g**), the 7.72 µm HiP-CT dataset (**b**,**e**,**h**) and the 100 µm “Edlow Brain” [3] dataset from doi:10.18112/openneuro.ds002179.v1.1.0 (**c**,**f**,**i**). **a**,**b**,**c**, Full coronal sections of the three datasets. **d**,**e**,**f**, Insets around the whole lentiform nucleus. **g**,**h**,**i**, Insets within the lentiform nucleus.

**Fig. A5.**
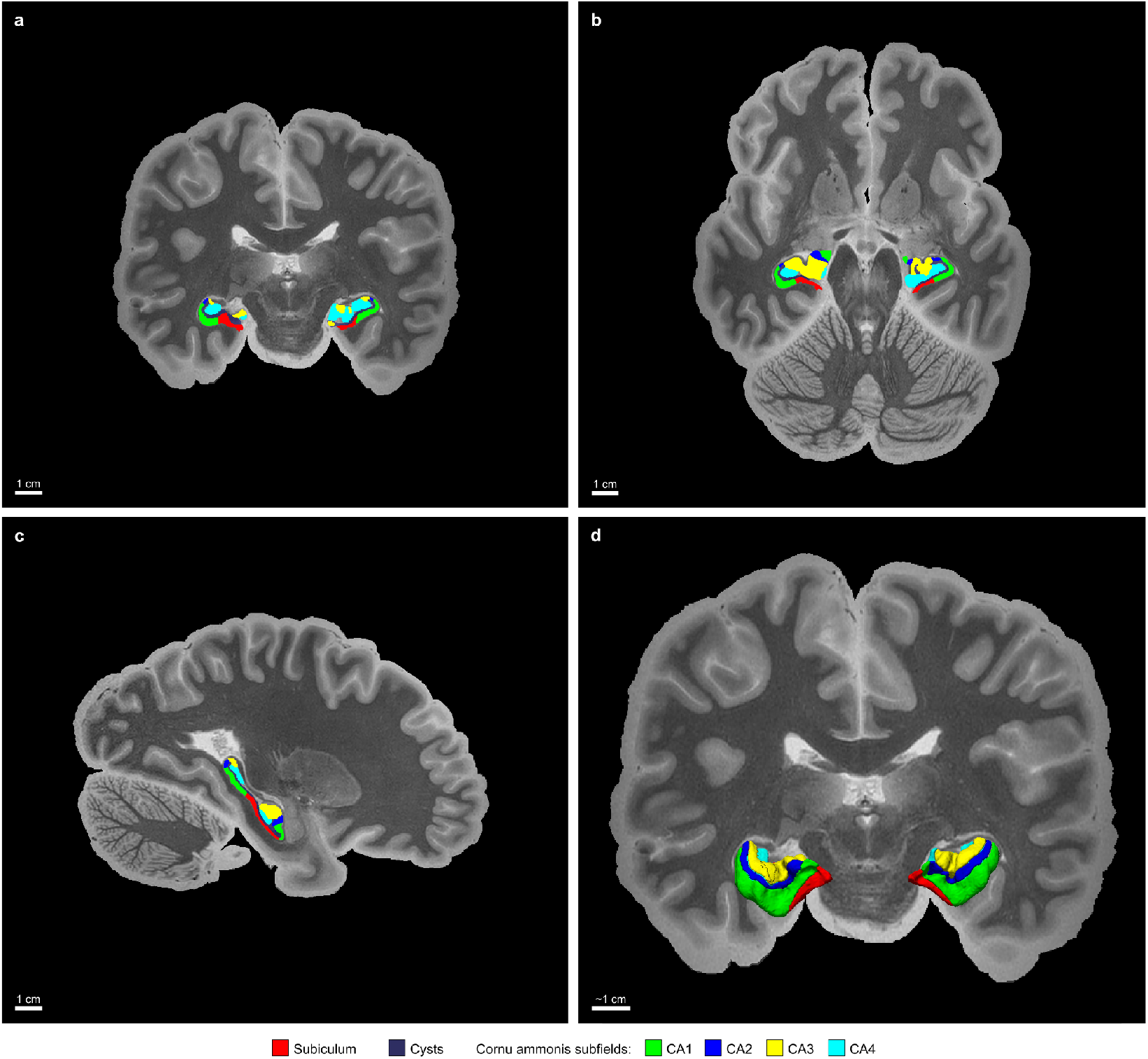
Overlay of the HippUnfold U-Net output on the 3 T *T*_2_-weighted (in-plane resolution: 0.255 mm; slice thickness: 0.257 mm). **a**, Coronal plane. **b**, Axial plane. **c**, Sagittal plane. **d**, 3D rendering of the labels over the coronal plane.

**Fig. A6.**
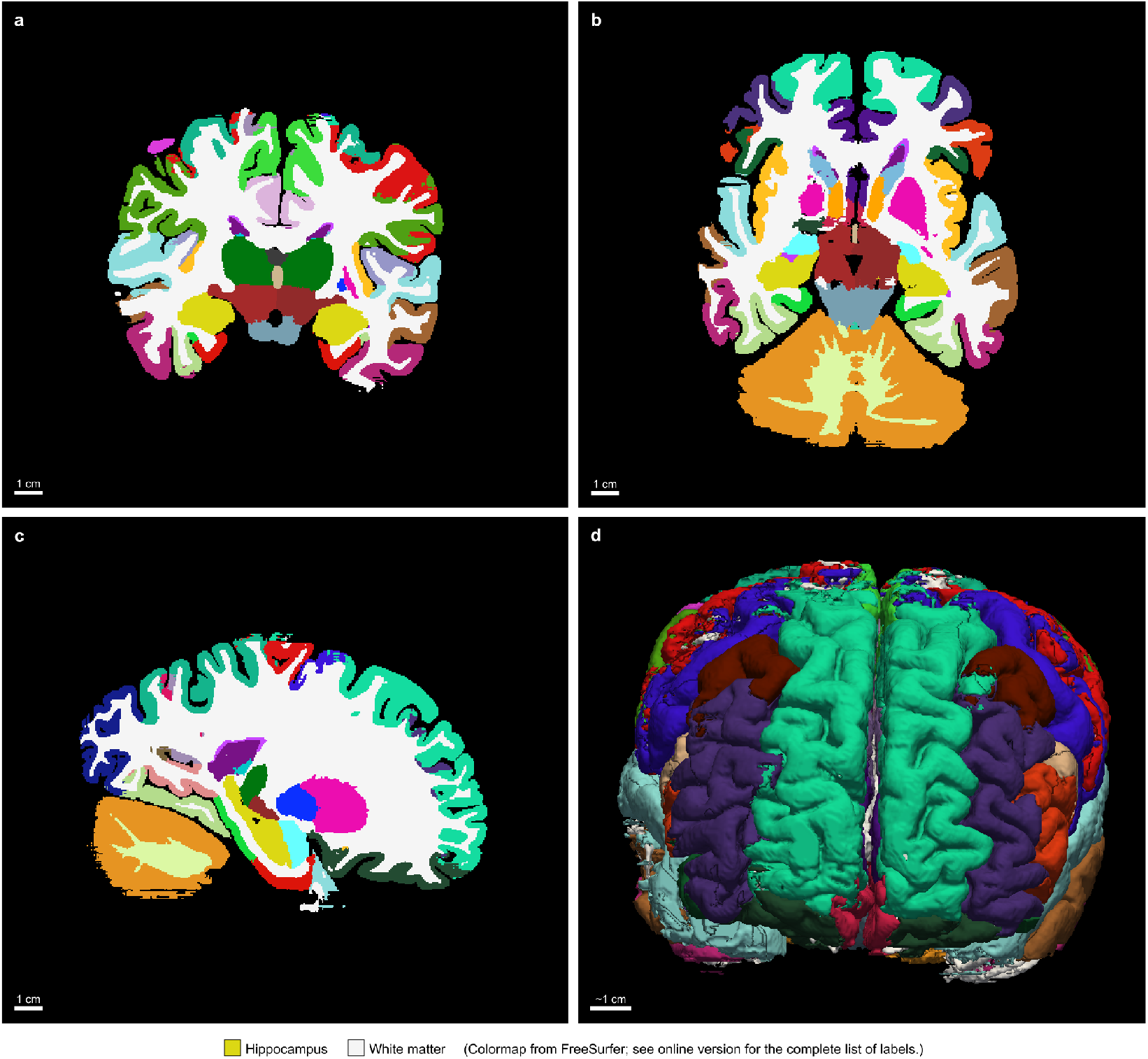
Initial output from the FastSurfer segmentation algorithm on the binned “Extremely Brilliant Brain” (EBB) dataset (isotropic voxel size of 494.08 µm). **a**, Coronal plane. **b**, Axial plane. **c**, Sagittal plane. **d**, 3D rendering of the labels. A few errors can be spotted within the segmentation. The segmentation of the left hippocampus was then manually refined, as shown in Figure 4. The colormap (or lookup table) is from the FreeSurfer software suite.

**Table A1.**
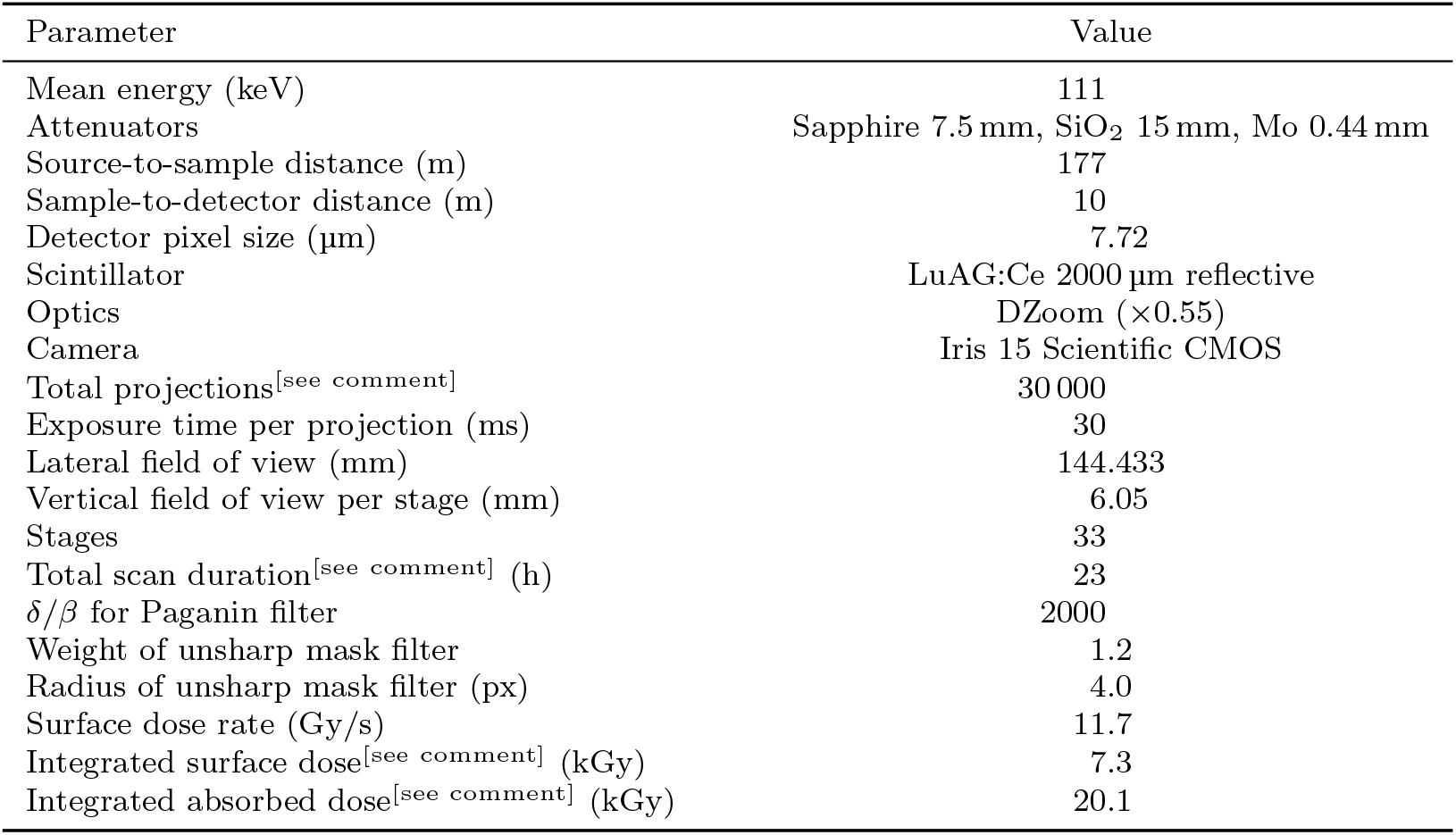
Acquisition and reconstruction parameters for HiP-CT. As mentioned in the main text, the quarter-acquisition scanning technique has produced two sets of 30 000 projections which were stitched into 30 000 projections. The total scan duration is the exposure time per projection multiplied by the number of projections, multiplied by two (quarter-acquisition), along with the motor overheads, movement duration and reference scans. The integrated surface dose corresponds to the surface dose rate multiplied by the total scan duration. The integrated absorbed dose takes into account the transmission of the X rays along the whole volume.

**Table A2.** Available scales of the dataset in the OME-Zarr. The binning of 1 is the native resolution and it gives the most details, making it relevant for numerical analysis; its size on disk as uncompressed raw 16-bit data is 15 063 GB ≈13.70 TiB. Binning the data enables the use of different software or hardware with limited resources. Parcellation was generated at 494.08 µm with FastSurfer on the windowed 8-bit volume. Masking was performed at 123.52 µm with 3D Slicer.

| Binning | Voxel size ($\mu\text{m}$) | Dimensions (voxels) | Size (GB) | Usecase |
| --- | --- | --- | --- | --- |
| 1 | 7.72 | 18 709 × 18 709 × 21 517 | 15 063 | Analysis |
| 2 | 15.44 | 9355 × 9355 × 10 759 | 1883 | |
| 4 | 30.88 | 4678 × 4678 × 5380 | 235 | |
| 8 | 61.76 | 2339 × 2339 × 2690 | 29.43 | High-resolution cinematic rendering |
| 16 | 123.52 | 1170 × 1170 × 1345 | 3.682 | Manual masking of background |
| 32 | 247.04 | 585 × 585 × 673 | 0.460 | Closest match to paired *ex vivo* $T_2$ MRI |
| 64 | 494.08 | 293 × 293 × 337 | 0.058 | Automatic parcellation with FastSurfer |
| 128 | 988.16 | 147 × 147 × 169 | 0.007 | Similar to clinical $T_1$ MRI |
| 256 | 1976.32 | 74 × 74 × 85 | 0.001 | |

